# Age- and sex- dependent changes in locus coeruleus physiology and anxiety-like behavior in response to acute stress

**DOI:** 10.1101/2020.11.10.377275

**Authors:** Olga Borodovitsyna, Daniel J Chandler

## Abstract

Adolescence is a critical period of development with increased sensitivity toward psychological stressors. Many psychiatric conditions emerge during adolescence and animal studies have shown that that acute stress has long-term effects on hypothalamic pituitary adrenal axis function and behavior. We recently demonstrated that acute stress produces long-term electrophysiological changes in locus coeruleus and long-lasting anxiety-like behavior in adolescent male rats. Based on prior reports of increased stress sensitivity during adolescence and increased sensitivity of female locus coeruleus toward corticotropin releasing factor, we hypothesized that the same acute stressor would cause different behavioral and physiological responses in adolescent female and adult male rats one week after stressor exposure. In this study, we assessed age and sex differences in how an acute psychological stressor affects corticosterone release, anxiety-like behavior, and locus coeruleus physiology at short- and long-term intervals. All groups of animals responded to stress with elevated corticosterone levels at the acute time point. One week after stressor exposure, adolescent females showed decreased firing of locus coeruleus neurons upon current injection and increased exploratory behavior compared to controls. The results were in direct contrast to changes observed in adolescent males, which showed increased anxiety-like behavior and increased spontaneous and induced firing locus coeruleus neurons a week after stressor exposure. Adult males were both behaviorally and electrophysiologically resilient to the long-term effects of acute stress. Therefore, there may be a normal developmental trajectory for locus coeruleus neurons which promotes stress resilience in adults, but stressor exposure during adolescence perturbs their function. Furthermore, while locus coeruleus neurons are more sensitive to stressor exposure during adolescence, the effect varies between adolescent males and females. These findings suggest that endocrine, behavioral, and physiological responses to stress vary among animals of different age and sex, and therefore these variables should be taken into account when selecting models and designing experiments to investigate the effects of stress. These differences in animals may also allude to age and sex differences in the prevalence of various psychiatric illnesses within the human population.

## 1. Introduction

Adolescence is an important period of transition between childhood and adulthood and is a sensitive period for stressor exposure (van Dijken et al. 1993). Neuroendocrine development in adolescence is characterized by maturation of the central nervous system (CNS) and development of hypothalamic pituitary adrenal (HPA) axis. A surge of sexual hormones during puberty makes the adolescent HPA axis more susceptible to negative effects of stress. For example, it takes almost twice as long for adolescent rats to return to the normal baseline level of the stress hormone corticosterone, compared to adults, during recovery after stress (Romeo et al. 2004, Romeo et al. 2004). Behaviorally, adolescence in humans is characterized by a pursuit of independence, changing social status, and increased risk-taking behavior. These changes are paralleled in rodents by increased play behavior and social interactions, and greater exploration of novel environments (Spear 2000). It has been reported that peripubertal stress leads to increased risk-taking behavior and decreased anxiety-like behavior in late adolescence (Toledo-Rodriguez and Sandi 2011). Increased behavioral and emotional susceptibility to stress and risk-taking behavior in adolescence could be explained in part by faster development of limbic structures, including the central nucleus of the amygdala, compared to prefrontal cortex (PFC) (Arain et al. 2013). This disparity in the rate of development of limbic structures could make PFC even more vulnerable to the negative effects of stress (Arnsten 2009) during adolescence. There have also been inconsistent reports of the effects of stress on behavior, cognitive function, and HPA axis responsiveness depending on the gender, sex, and type of stress used (McCormick et al. 2010). Rarely has more than one treatment group been included in studies, so additional research is necessary to compare the stress effects across different ages and sexes.

A major brain nucleus involved in the modulation of the central stress response is the locus coeruleus/norepinephrine (LC/NE) system. It is activated by stress through release of corticotropin releasing factor (CRF) which interacts with corticotropin releasing factor receptor 1 (CRFR1). This increases tonic firing of LC neurons and promotes forebrain release of NE (Valentino et al. 1983, Valentino et al. 1992, Jedema and Grace 2004). While there is an established relationship between LC discharge rates and behavioral indices of arousal and anxiety-like behavior, the long-term effects of stress on LC, and its relationship to behavior, is less clear. The transition from adolescence into adulthood is characterized by a number of changes in the LC/NE system: decreased transporter content in PFC (Bradshaw et al. 2016), age-related decline in LC spontaneous cell firing (Olpe and Steinmann 1982) and increased capacity for the recovery and adaptation in response to social stress (Zitnik et al. 2016). Accordingly, it is worthwhile to investigate if LC might mediate chronic changes in both physiology and behavior induced by an acute stressor. Indeed, studies have shown that behavioral and physiological changes occur in adulthood after adolescent stress (Yohn and Blendy 2017). Postnatal environmental changes are also known to impact the developing brain (Kolb and Gibb 2011) and thus stressor exposure in adolescence may chronically alter LC in such a way that changes behavior in later life.

We have previously shown that a single acute stressor exposure (fifteen minutes combined predator odor and physical restraint) is sufficient to produce behavioral and LC electrophysiological changes a week after stress (Borodovitsyna et al. 2018) in adolescent male rats. The study included only male rats and it is known that the LC is sexually dimorphic in both morphology and stress responsiveness (Bangasser, Wiersielis, and Khantsis 2016). Specifically, there is evidence for denser afferentation from limbic areas in female LC, prolonged activity of CRF receptor activation on LC neurons as a result of slower receptor internalization, and presynaptic modulation of NE release with increased estrogen-dependent release and decreased degradation of NE (Bangasser, Wiersielis, and Khantsis 2016). Sex-differential expression of more than 100 genes was also identified in adult mouse LC (Mulvey et al. 2018). Based on these findings, a more comprehensive analysis of how age, sex, and stress interact to modulate anxiety-like behavior is necessary. Therefore, in this study, both adolescent males and females, as well as adult male rats are included. Here we describe age- and sex-specific electrophysiological changes of LC neurons and anxiety-like behavior in response to stress in multiple assays at different time points. We further assess the HPA axis response based on serum corticosterone levels. Together endocrine, physiological and behavioral responses to stressor exposure among animals of both sexes and different ages will help to create a more refined picture of the effects of acute stressor exposure on the LC/NE system.

## 2. Material and methods

### 2.1: Subjects

Adolescent male (30 PND), adolescent female (35 PND) and adult (77 PND) male Sprague Dawley rats (Taconic Farms), were housed two to three per cage on a 12 h reverse light schedule (lights on at 9:00pm) with access to standard rat chow and water *ad libitum*. Adolescent females we chosen to be 5 days older to match the weight of adolescent males. All effots were made to minimize animal suffering and numbers. All animal protocols were approved by the Rowan University Institutional Animal Care and Use Committee and were conducted in accordance with National Institutes of Health *Guide for the Care and Use of Laboratory Animals*.

### 2.2: Stressor Exposure

Essentially as previously described (Borodovitsyna, Flamini et al. 2018). Briefly, to induce acute stress, rats were placed in a rodent restrainer (Harvard Apparatus) for 15 minutes. The restrainer was then placed inside of the plastic chamber. Delivery of the predator odor 2,4,5-trimethylthiazole (TMT, Sigma-Aldrich) was achieved by connecting the chamber via silicone tubing to an aquarium pump. Odor was delivered continuously during the 15 minutes by turning on the aquarium pump, which forced odor into the sealed odor exposure chamber. Control animals were placed in an identical bell chamber for fifteen minutes, but they were not restrained and no odor was delivered.

### 2.3: Serum corticosterone

In some of the animals used for behavioral and electrophysiological experiments blood (0.2-0.4 mL) was collected from the saphenous vein immediately before the onset of stress or control conditions (labeled as “baseline”) and 35 min after the onset (labeled as “35 min”). In some subjects who were studied for long-term effects, additionally to the baseline and acute collections, blood sample (0.2-0.4 mL) was collected from the saphenous vein after behavioral testing one week after control or stress conditions and was labeled as “1 week”. In all cases, blood was collected per the protocol for frequent blood collection from lateral saphenous vein in unanesthetized animals (Beeton, Garcia, and Chandy 2007; Parasuraman, Raveendran, and Kesavan 2010). Blood was collected into 1.5mL Eppendorf tubes and left for at least 10 min to coagulate, and then centrifuged at 3000 r.p.m. for 4 min. Serum was kept at −80°C prior to analysis using a Corticosterone Enzyme Immunoassay kit from Enzo (ADI-900-097). Results were calculated using Four Parameter Logistic Curve fit in R-Studio.

### 2.4: Behavioral tests

#### 2.4.1: Elevated plus maze

Essentially as previously described (Borodovitsyna, Flamini et al. 2018). Immediately after exposure to stress or control conditions, rats were placed in the center of an elevated plu s maze (EPM). Rats were allowed to explore the maze for 10 minutes, during which their activity was filmed with an infrared camera. At the conclusion of each test, rats were returned to their home cage for a week. Behavior was scored using AnyMaze behavioral tracking software (Stoelting).

#### 2.4.2: Open field test

Essentially as previously described (Borodovitsyna, Flamini et al. 2018). One week after testing in the EPM, rats were placed in the center of an open field test (OFT) apparatus. The OFT apparatus consisted of a 90cm × 90cm × 30cm black plexiglass box. Rats were allowed to explore the apparatus for 10 minutes, during which their activity was filmed with an infrared camera situated above the maze. At the conclusion of each test, rats were sacrificed for electrophysiological recordings. Behavior was scored using AnyMaze behavioral tracking software (Stoelting). Rats were tested in different mazes at the two time points to eliminate the possibility of habituation to any one test confounding anxiety-like behavior.

### 2.5: Whole cell patch clamp electrophysiology

#### 2.5.1: Brain slice preparation

Essentially as previously described (Borodovitsyna, Flamini et al. 2018). Briefly, rats were deeply anesthetized with an intraperitoneal injection of Euthasol (100mg/kg, Virbac) and transcardially perfused with ice cold oxygenated artificial cerebrospinal fluid (aCSF). Rats were then rapidly decapitated and the skull was removed so that gross coronal cuts could be made at the level of the medulla and the pineal gland. The blocked brain was submerged in the sucrose-aCSF for 1-2 minutes after which it was transferred to a piece of filter paper. The dorsal aspect of the brain was cut at 200μM thick horizontal sections using Compresstome VF-300-0Z tissue slicer. Sections containing LC were transferred into aCSF continuously bubbled with 95% O_2_/5% CO_2_ and maintained at 35.5°C. After 1 h, the holding incubator was maintained at room temperature.

#### 2.5.2: Electrophysiological recordings

Essentially as previously described (Borodovitsyna, Flamini et al. 2018). Slices were individually transferred to a recording chamber which was continuously superfused with oxygenated aCSF and maintained at 37°C by a Warner Instrument Corporation in-line heater (model 60-01013). LC was visualized as a semi-translucent crescent-shaped region located lateral to the fourth ventricle at 5X magnification using an Olympus BX51WI fixed-stage upright microscope with differential interference contrast and an infrared filter. Individual LC neurons were visualized with a 40X immersion lens and QImaging Rolera Bolt camera using QCapture Pro software. Neurons were approached with sharp glass electrodes (resistance = 5-10MΩ) controlled with Sutter MPC-200 manipulators. Electrodes were filled with intracellular solution of the following composition, in mM: KCl 20, K-gluconate 120, MgCl_2_ 2, EGTA 0.2, HEPES 10, Na_2_ATP 2, biocytin. After a GΩ seal was established between the pipette and neuronal membrane, the membrane was ruptured and whole-cell recordings were obtained. To assess membrane properties in current clamp mode, spontaneous activity was recorded for 60s after stabilization of cell firing and the average firing rate was calculated. Cells were then subjected to a series of increasing current steps from −250pA to 300pA with 50pA intervals between sweeps, and the input resistance and number of action potentials fired in response to each level of current was determined.

### 2.6: Data analysis

Electrophysiological data were recorded using Molecular Devices ClampEx 10.2 and MultiClamp 700B acquisition software and analyzed with ClampFit 10.2. Behavioral data were acquired and analyzed using AnyMaze behavioral tracking software (Stoelting). Statistical analyses were performed with R-studio Version 1.1.453. Data was tested for normality using Shapiro-Wilk test. Normally distributed data were tested for equal variances of distribution, and for the data with equal variances two-sided t-test was performed, while if variances were unequal Welch modification of t-test was conducted. Wilcoxon Rank Sum test (Mann Whitney U test) was used for non-normally distributed data sets. Pearson correlation coefficient was calculated. Statistical significance was set with alpha = 0.05. Error bars in all figures are presented as mean ± SEM unless otherwise is indicated.

## 3. Results

### 3.1: Adolescent females

To demonstrate effects of acute stress on adolescent female rats the study was performed according to the experimental timeline shown in Figure 1. Serum corticosterone levels from female rats are shown in Figure 2A. Acute exposure to TMT and restraint stress significantly increased corticosterone levels 35 min after the beginning of stress exposure compared to control animals (t = −4.4089, df = 11, p = 0.001). No significant differences in corticosterone levels between stress and control rats were identified at either baseline or one week measurements.

**Figure 1.**
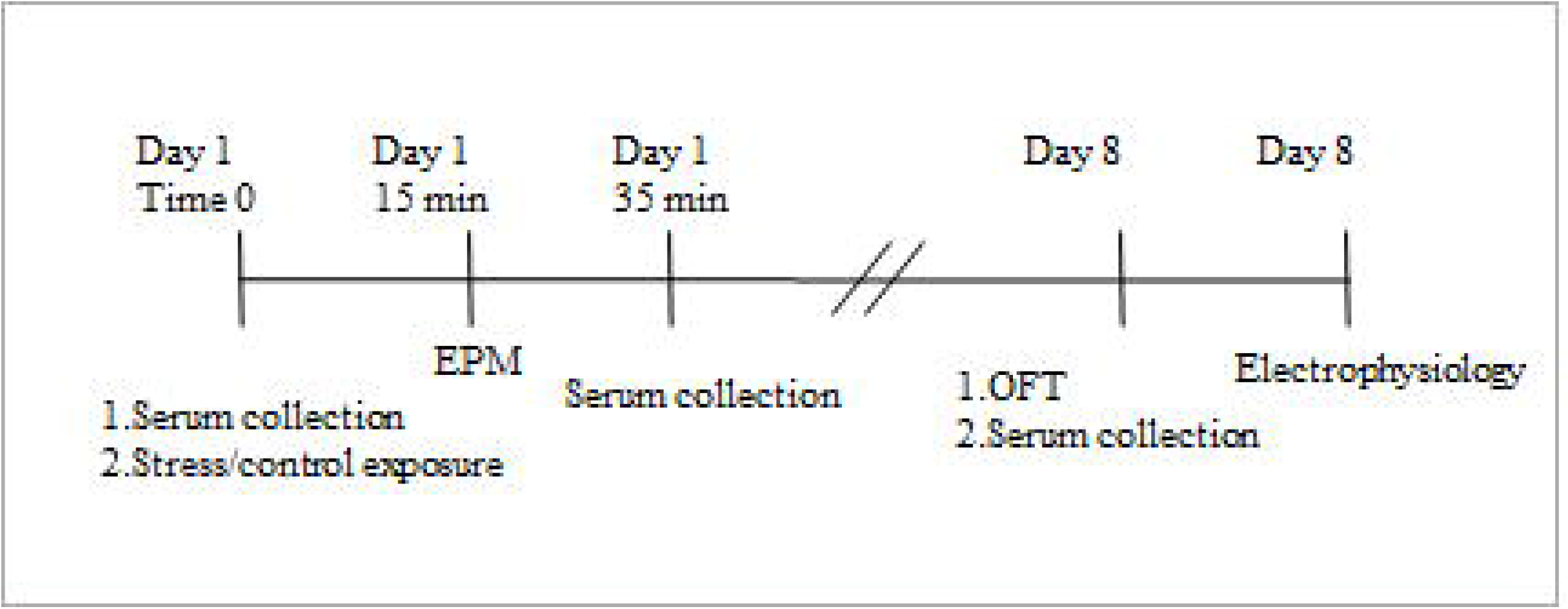
Experimental timeline.

**Figure 2.**
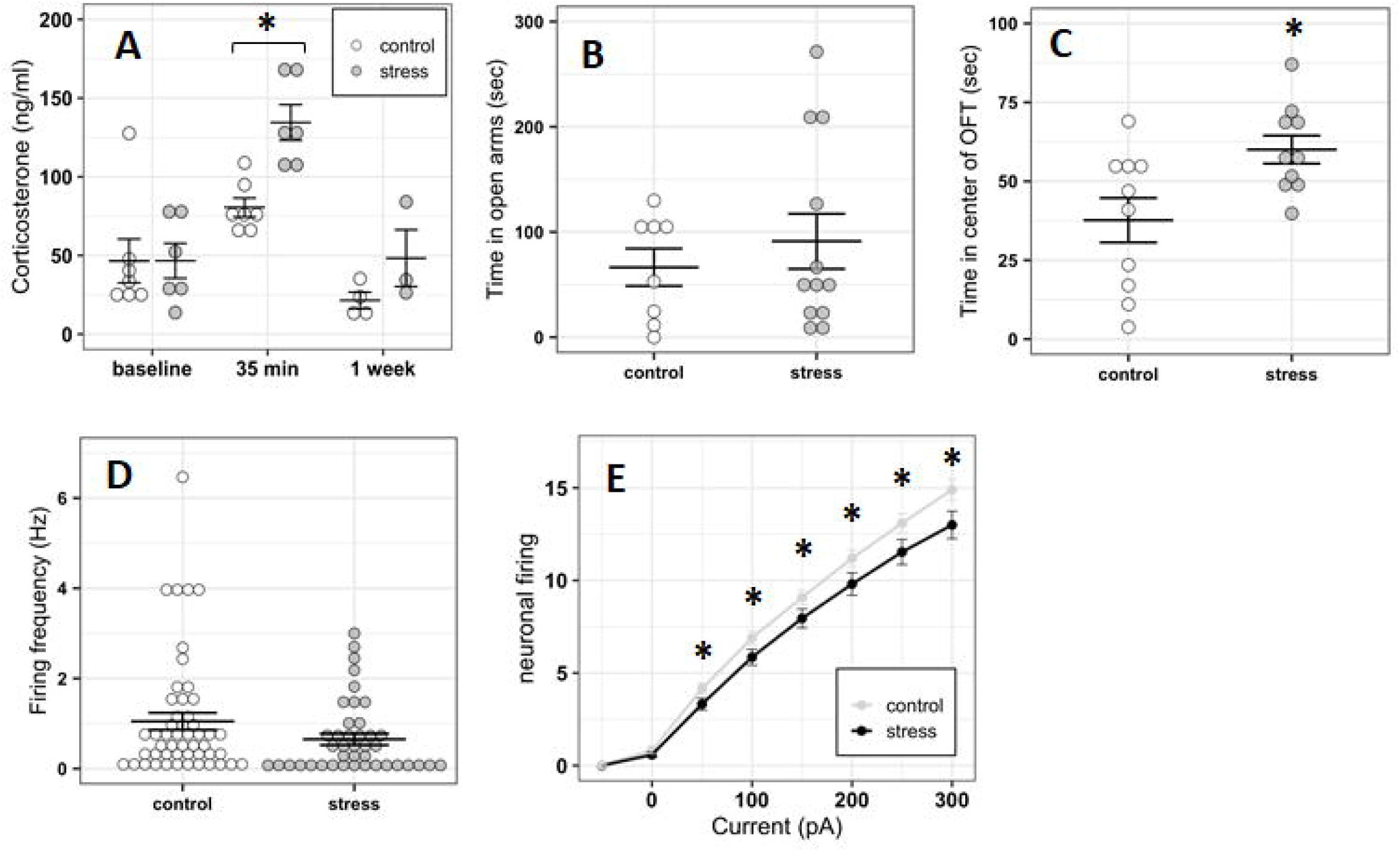
Stress responses in adolescent female rats. (A) Serum corticosterone prior to (control=7 animals, stress=6 animals), 35 min (control=7 animals, stress=6 animals), and 1 week (control=4 animals, stress=3 animals) after stress or control conditions. (B) Time spent in open arms of elevated plus maze immediately after stress (n=12) or control (n=8). (C) Time spent in the center zone of open field test a week after stress (n=10) or control (n=10). (D) Spontaneous firing of LC neurons a week after stress (n=41 cells from 7 animals) or control (n=50 cells from 8 animals). (E) Induced firing upon the current injection with 50 pA step from −50pA to 300 pA.

Anxiety-like behavior was assessed immediately after (Figure 2B) and one week after (Figure 2C) stress in the EPM and OFT, respectively. Time spent in the open arms and time spent in the center of open field were used as measurements of anxiety-like behavior. We did not observe any statistical difference in time spent in the open arms in the EPM between control and stress rats. However, stressed female rats spent significantly more time in the center of the OFT a week after stress (t = −2.6908, df = 15.181, p = 0.017).

There was a trend towards but no significant effect of stress on spontaneous firing rate of female LC neurons a week after stress (W = 1242.5, p = 0.08225; Figure 2D). There was a statistically significant effect of stress on evoked LC firing in response to current injection for all current steps beyond 50 pA (Figure2E): 50 pA (W = 1308.5, p-value = 0.03659), 100 pA (W = 1353, p-value = 0.01484), 150 pA (W = 1303.5, p-value = 0.04159), 200pA (W = 1328, p-value = 0.02584), 250pA (W = 1343, p-value = 0.01905), 300 pA current injection (W = 1368, p-value = 0.01108), such that cells from stressed rats fired less frequently in response to current injection than those from control animals.

### 3.2: Adolescent males

Adolescent male rats were subject to the same experimental timeline as adolescent females (Figure 1). Corticosterone values were significantly higher in animals exposed to the stressor at the 35 min time point compared to controls (t = −2.6893, df = 27, p = 0.012 (Figure 3A). There were no differences between groups at the baseline or one week time points.

**Figure 3.**
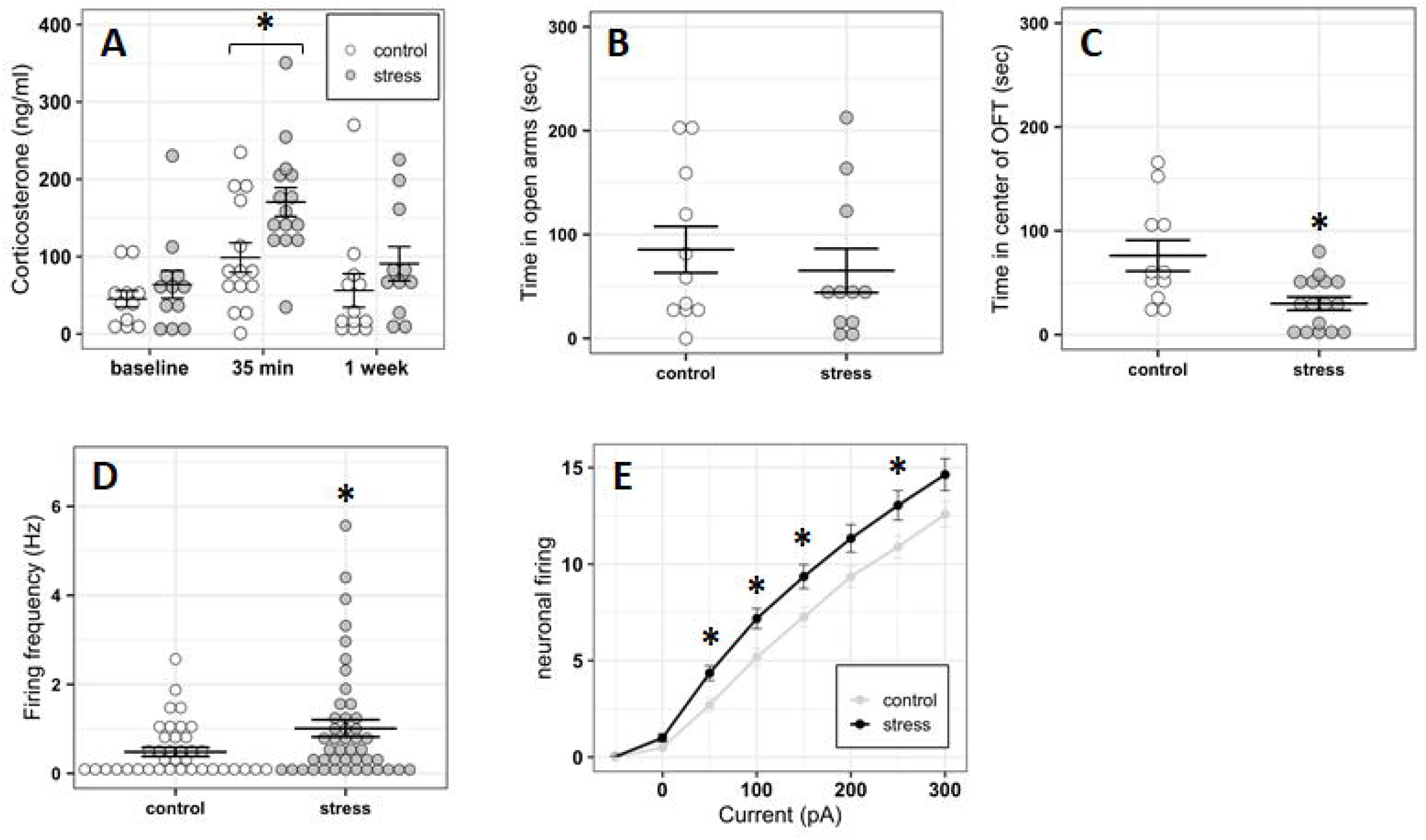
Stress response in adolescent male rats (A) Serum corticosterone prior to (control=11 animals, stress=12 animals), 35 min (control=14 animals, stress=15 animals), and 1 week (control=12 animals, stress=11 animals) after stress or control conditions. (B) Time spent in open arms of elevated plus maze immediately after stress (n=11) or control (n=11 animals). (C) Time spent in the center zone of open field test a week after stress (n=16), control (n=11). (D) Spontaneous firing of LC neurons a week after stress (n=45 cells from 8 animals), or control (n=38 cells from 8 animals). (E) Induced firing upon the current injection with 50 pA step from - 50pA to 300 pA.

We did not observe a difference in the behavior in EPM immediately after stress (Figure 3B). However, stressed adolescent male rats spent significantly less time in the center of the OFT a week after stress (Figure 3C, W = 141.5, p = 0.009). Consistent with our previously published results, LC neurons from stressed adolescent male rats fired more frequently compared to those from control animals (Figure 3D, W = 620, p = 0.031). The induced firing of LC neurons in response to current injection was significantly different between groups at 50 pA (W = 558.5, p-value = 0.006155), 100pA (W = 586.5, p-value = 0.01356), 150pA (W = 590.5, p-value = 0.0152) and 250 pA current injection (W = 624.5, p-value = 0.04877, Figure 3E). In contrast to adolescent female rats, neurons from stressed animals fired more frequently in response to current injection compared to those from control animals (Figure 2E).

### 3.3: Adult males

Adult male rats were subject to the same experimental timeline as the other groups (Figure 1). Corticosterone levels were significantly higher in the stressor-exposed rats at the 35 min time point relative to controls (t = −2.1101, df = 20, p = 0.048, Figure 4A). No such differences were observed between groups at the baseline or one week time points.

**Figure 4.**
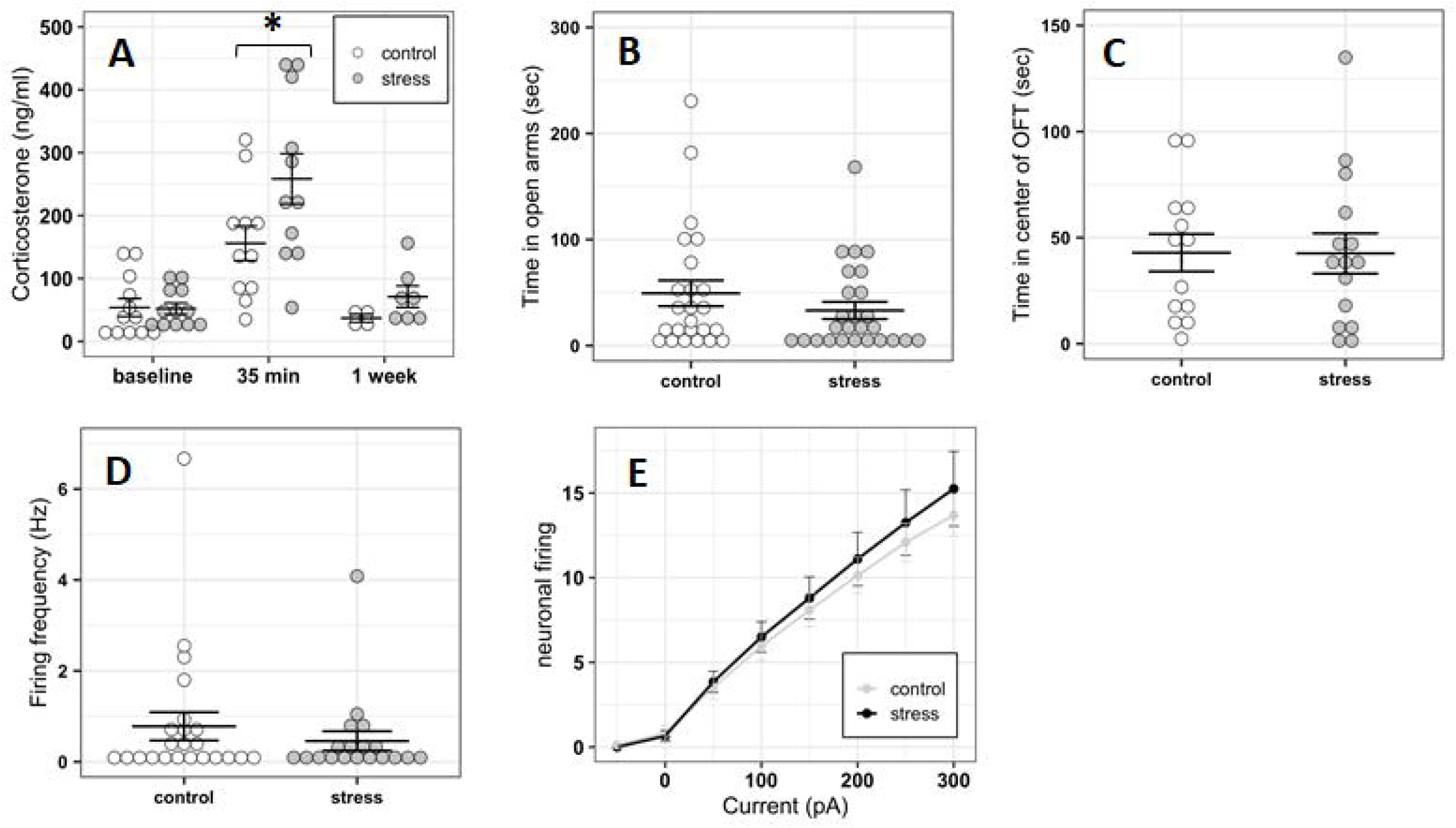
Stress response in adult male rats (A) Serum corticosterone prior to (control=12 animals, stress=12 animals), 35 min (control=11 animals, stress=11 animals), and 1 week (control=4 animals, stress=7 animals) after stress or control conditions. (B) Time spent in open arms of elevated plus maze immediately after stress (n=26) or control (n=24). (C) Time spent in the center zone of open field test a week after stress (n=12) or control (n=16). (D) Spontaneous firing of LC neurons a week after stress (n=20 cells from 3 animals) or control (n=23 cells from 3 animals). (E) Induced firing upon the current injection with 50 pA step from −50pA to 300 pA.

Anxiety-like behavior of adult male rats was unaffected by stressor exposure in both the EPM at the acute time point and the OFT at the one week time point (Figures 4B&C). Accordingly, spontaneous and induced LC neuronal firing rates did not differ between stress and control groups a week after stress (Figures 4D&E).

### 3.4: Correlation analysis

Because of a clearer relationship between stressor exposure and anxiety-like behavior in adolescent male rats, we sought to explore how corticosterone levels in this group at the 35 min and one week timepoints in this group related to behavior in both the EPM and OFT. Correlation mapping of data from all adolescent males (both stress and control) shows that both time spent in the open arms of EPM (R^2^=0.2832, p=0.003, Figure 5B) and distance traveled in the EPM (R^2^=0.42, p= 0.0001, Figure 5A) are significantly negatively correlated with corticosterone levels at the 35 min time point. Similarly, considering all adolescent male rats regardless of treatment, there were significant negative correlations between corticosterone at the one week time point and distance traveled in the OFT (R^2^= 0.3054, p=0.006, Figure 5C) and corticosterone at the one week time point and number of entries to the center of the OFT (R^2^=0.2144, p=0.026, Figure 5D). No correlations between firing frequency and behavior were identified in adolescent males. No significant correlations were identified in adolescent females or adult males.

**Figure 5.**
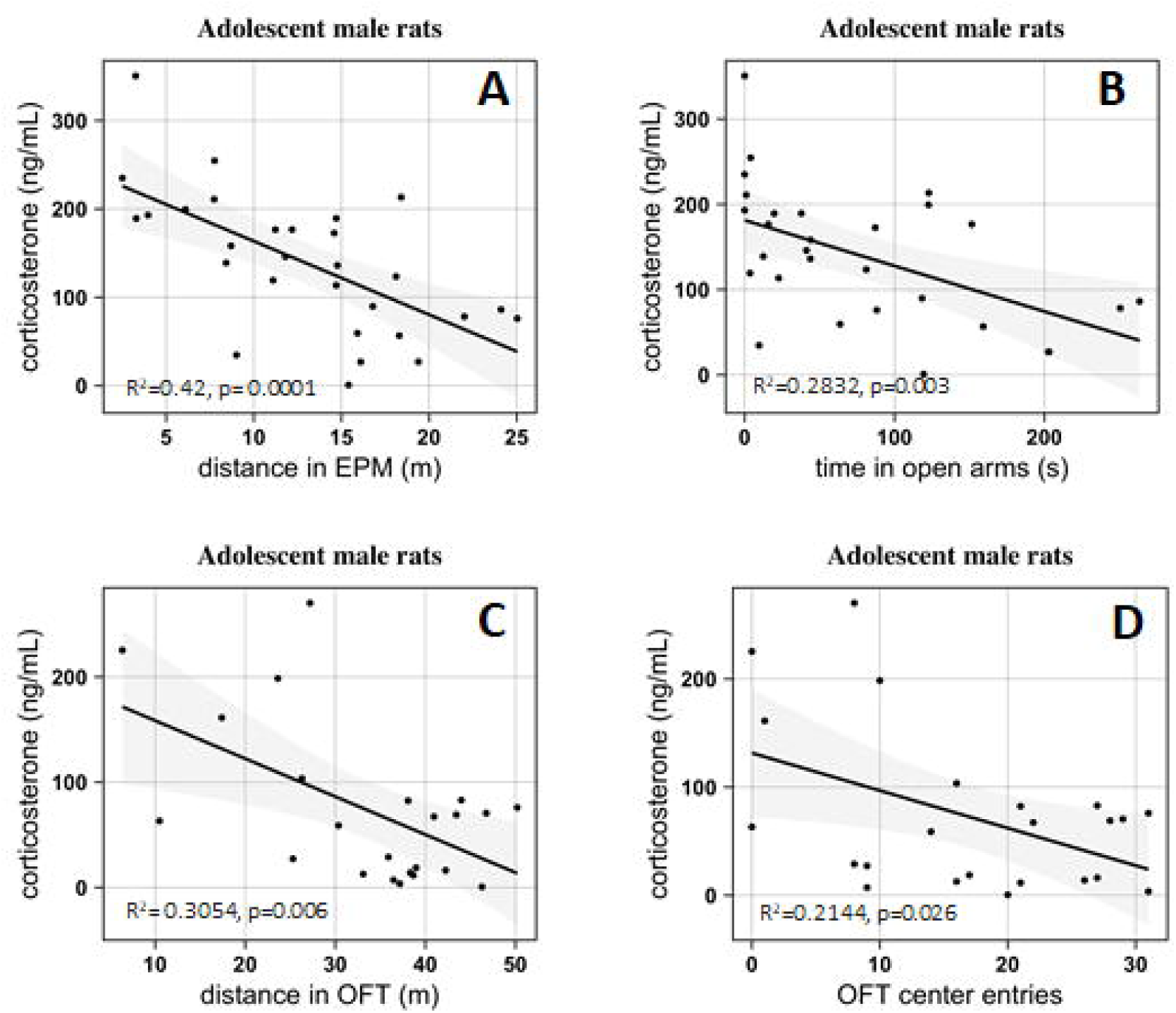
Anxiety-like behavior significantly correlated with corticosterone levels in adolescent male rats. Correlations between corticosterone levels immediately after stress/control exposure and distance travelled in elevated plus maze (A), between corticosterone levels immediately after stress/control exposure and time spent in open arms of elevated plus maze (B), between corticosterone levels one week after stress/control exposure and distance travelled in the open field test (C), and between corticosterone levels one week after stress/control exposure and number of entrances to the center zone of open field test (D). Shaded region corresponds to 95% confidence interval.

## 4. Discussion

In this study we demonstrated how acute stress differentially affects hormonal, electrophysiological and behavioral responses in rats of varying age and sex. We previously demonstrated the effect of combined restraint and predator odor stress (Borodovitsyna, Flamini et al. 2018) on adolescent male LC physiology and anxiety-like behavior. Here, we extend these findings to show how the same stressor affects HPA axis activation, LC physiology and behavior in adolescent female, adult male and adolescent male rats.

The HPA axis is the central regulator of stress response. Its activation results in the release of steroidal hormones by the adrenal glands, which leads to energy metabolism mobilization in order to adapt to an adverse situation (Selye 1975). We observed significantly elevated levels of corticosterone 35 min after stress that also were similar in all animal groups (Figures 2A, 3A& 4A). This confirms that the stressor used in this study was sufficient to produce an endocrine response in all groups. The slight but statistically non-significant increase in corticosterone in control animals at 35 min can likely be attributed to the brief restraint and pain associated with unanesthetized blood drawing. A week after stress, corticosterone concentration returned to baseline levels in all groups, regardless of treatment groups (Figures 2A, 3A& 4A), which indicates that the stressor does not result in chronic dysregulation of the HPA axis. HPA axis activation starts with the release of corticotropin releasing hormone (CRH) by the paraventricular nucleus (PVN) of hypothalamus. While it is known that there is a CRF-containing projection from hypothalamus to LC (Reyes et al. 2005), it is unclear if the changes in LC physiology we identified in adolescent males and females relate directly to activation of the HPA axis. This is due to the fact that we did not detect a direct relationship between serum corticosterone levels and LC firing rates and only were able to find correlations between HPA axis activation and behavior (Figure 5). Although the HPA axis and LC/NE systems are both activated during stress, observations from this study suggest that they may operate independently of one another and not in a sequential or ordered fashion. It is also interesting to note that although the hormonal response to stress is similar in adolescent males, adolescent females and adult males, the adolescent male LC is particularly susceptible to stress-induced physiological changes, and that behavioral adaptations are different in response to stress among these groups. Specifically, adolescent female rats showed a significant stress-induced increase in time spent in the center of the OFT, while males showed a significant decrease. These findings indicate that there are behavioral long-term adaptations in response to stress, but they vary according to the sex of the animal. It is difficult to assess whether this corresponds directly to a stress-induced pro-exploratory and/or anxiolytic effect in females, or if anxiety-like behavior in the open field manifests differently in males and females. Interestingly, however, these findings parallel the stress-induced physiological changes we identified in LC. Specifically, females showed increased time in the center of the open field and decreased neuronal excitability, while males showed decreased time in the center of the open field and increased spontaneous and evoked LC discharge. Given the well-established relationship between LC discharge rates and anxiety-like behavior, it can at least be postulated that stressor exposure produces an anxiolytic effect in females one week later.

These data match well with findings from other studies which have also shown that adolescent female and male rats differ in their stress response. For example, female Wistar rats have been shown to display increased exploratory behavior and reduced anxiety-like behavior later in adulthood after either lipopolysaccharide-induced inflammatory stress or acute restraint stress during adolescence, while no changes were observed in male rats (Ariza Traslavina et al. 2014). Adolescent female Long-Evans rats were also shown to have decreased anxiety-like behavior after social stress without any effect on males (McCormick et al. 2008). On the other hand, it has been reported that peripubertal stress leads to increased risk-taking behavior and novelty seeking in Wistar Han male and female rats with more pronounced effect in females (Toledo-Rodriguez and Sandi 2011). Our findings from adolescent female and male rats are also consistent with another recent publication (Lovelock and Deak 2019), which used acute footshock stress during adolescence (30 PND) and reported long-term increased anxiety-like behavior in males but decreased anxiety-like behavior in females in adulthood (70 PND) (Lovelock and Deak 2019).

The fact that there is a disparity in reports of sex differences in stress responses may be explained in part by the nature and the duration of the stress (Ariza Traslavina, de Oliveira et al. 2014). While the existing body of literature on this topic reviews the behavioral effects of stress, there are few studies of sex differences in stress responsiveness of the LC/NE system in adolescent rats, despite its sexually dimorphic nature and a clear role in mediating arousal, vigilance, and anxiety-like behavior. Therefore, we assessed electrophysiological characteristics of LC neurons using whole-cell patch clamp electrophysiology. We found that LC neurons from stressed adolescent females have decreased evoked activity compared to control LC neurons. This was a surprising finding because the female LC/NE system is known to be more susceptible to CRF, and has been demonstrated to be more sensitive to both acute hypotensive and swim stress (Curtis et al. 2006, Bangasser et al. 2010, Bangasser et al. 2016). Such decreased excitability of female LC neurons may be due to possible long-term effects of CRF signaling which alters gene expression and changes in ion channel trafficking (Borodovitsyna et al. 2018). Additionally, a recent genetic analysis (Mulvey, B., Bhatti, D. L. et al. 2018) demonstrated sex-differential expression of multiple genes in LC including elevated Prostaglandin E2 (PGE2) receptor Ptger3 (EP3) in female mice. The same study demonstrated that activation of EP3 with the agonist sulprostone led to decreased anxiety-like behavior in female mice after restraint stress and decreased firing in female LC neurons compared to males. Similar mechanisms might contribute to the long-term effects observed in our study.

Another relevant question that this study raises is why adolescent female rats show opposing to males changes in anxiety-like behavior and LC neuronal physiology in response to stress, while human females are generally more susceptible to stressor exposure and have almost twice as high of an incidence of psychiatric conditions such as posttraumatic stress disorder (PTSD) (Kessler et al. 1995), anxiety disorder (Bandelow and Michaelis 2015) and depressive disorder (Angst et al. 2002, Accortt et al. 2008). It has been demonstrated that repeated restrained stress decreases basolateral amygdala (BLA) neuronal firing and causes neuronal atrophy in the lateral nucleus of amygdala in females, while males show opposite changes in morphology and physiology (Blume et al. 2019). These data parallel our findings, and sex-specific changes in amygdala function contribute to the sex-specific stress-induced changes in LC physiology we have identified in adolescent rats. As mentioned above, it remains to be determined if the increased time in the center of the open field we identified in stressed females is an anxiolytic effect, or if coping strategies differ between males and females. For example, avoidant coping might contribute to peritraumatic and posttraumatic dissociation syndrome and peritraumatic distress are described more frequently among female patients (Boisclair Demarble et al. 2020). Therefore, impaired BLA function in response to repeated stress in female rats (Blume, Padival et al. 2019) combined with disrupted LC neuronal activity we observed in response to acute stress in female rats, might contribute to poor behavioral long-term adaptations in response to stress and higher risk of stress-associated psychiatric disorders in females. Amygdala is a major source of CRF for LC, and this pathway is responsible for LC activation and anxiety-like behavior formation during stress (McCall et al. 2015). A positive feedback activation loop was recently described from LC to BLA that contributes to anxiety-like behavior (McCall et al. 2017), indicating tight reciprocal control between these structures. Therefore, sexually dimorphic stress responses in one region may compound sexually dimorphic changes in the other.

In addition to the unique changes that we identified in LC physiology and behavior in male and female rats, there were also important differences between adolescent and adult male rats. Adolescent male rats showed increased anxiety-like behavior a week after stress that was associated with persistently increased LC neuronal spontaneous and evoked firing relative to control animals. This indicates that LC undergoes neuroadaptive changes in response to stressor exposure in adolescent male rats. Conversely, adult male rats did not demonstrate any changes in anxiety-like behavior or LC physiological characteristics in response to stressor exposure. It is important to acknowledge that we did not include adult females in this study. This is due to the long-term nature of our experiment which would require the inclusion of several different groups depending on the phase of the estrous cycle during stressor exposure and phase of the cycle during analysis and also existing data about estrous cycle effect on anxiety-like behavior in female rats (Mora et al. 1996). Future studies will explore how LC physiology and behavior change in response to stress according to estrous cycle phase. In humans, adolescence is a critical period of development when many psychiatric disorders first emerge due to structural and functional synaptic changes (Paus et al. 2008). Our data confirms that susceptibility of the adolescent LC/NE system to stress may contribute to the development of aberrant anxiety-like behavior. This is in contrast to the developed brain and LC/NE system observed in adults. Therefore, it is possible that during the transition from adolescence to adulthood, adaptations may occur in LC which confer resilience to stressor exposure at both the behavioral and physiological levels. While we did not specifically investigate in this study if LC properties differ between control adolescent and control adult male rats, others (Olpe and Steinmann 1982) have reported that there is an age-related decline in LC firing rates. Therefore, the normal developmental trajectory for LC neurons as they progress from and adolescence to adulthood may be a critical period when this development could be perturbed by stress. Our results are also consistent with findings from recent *in vivo* electrophysiology recordings after social stress (Zitnik, Curtis et al. 2016), in which adolescent male rats had elevated spontaneous firing of LC neurons while adult rats showed decreased firing. Another study similarly showed that social stress in early adolescence significantly increases LC spontaneous discharge relative to adult rats, and adult rats did not differ in LC tonic firing properties in response to social stress (Bingham et al. 2011).

## 5. Conclusions

The findings from these studies indicate that behavioral and LC neuronal responses to stress differ according to age and sex. These findings highlight the necessity to consider age and sex as important sources of variation when designing and performing experiments which investigate stress, behavior, and LC physiology. There are multiple factors involved in the development of psychiatric disorders such as anxiety disorders or PTSD, which include different brain areas and neurotransmitter systems, genetic predispositions and genetic polymorphisms, emotional and behavioral coping strategies. While LC is only one of many systems that contributes to behavioral state and the stress response, these findings help to clarify how it contributes to these phenomena among rats of different age and sex. These disparities may further allude to important differences between male and female humans of different ages in the prevalence, progression, and pathophysiology of various clinical psychiatric conditions.

## Declaration of interest

none.

## Acknowledgements

We wish to acknowledge Debra Bangasser, PhD for aid with experimental design, Matthew Flamini for aid with Four Parameter Logistic Curve modeling, and Brenna Duffy and John Tkaczynski for aid with manuscript preparation and editing.

## Funding

This work was supported by National Institutes of Mental Health [R56MH121918, 2019].

## Notes

### Competing Interest Statement

The authors have declared no competing interest.

## References

Accortt, E. E., M. P. Freeman and J. J. Allen, 2008. Women and major depressive disorder: clinical perspectives on causal pathways. J Womens Health (Larchmt) 17(10), 1583–1590.

Angst, J., A. Gamma, M. Gastpar, J. P. Lepine, J. Mendlewicz, A. Tylee and S. Depression Research in European Society, 2002. Gender differences in depression. Epidemiological findings from the European DEPRES I and II studies. Eur Arch Psychiatry Clin Neurosci 252(5), 201–209.

Arain, M., M. Haque, L. Johal, P. Mathur, W. Nel, A. Rais, R. Sandhu and S. Sharma, 2013. Maturation of the adolescent brain. Neuropsychiatr Dis Treat 9(449-461.

Ariza Traslavina, G. A., F. L. de Oliveira and C. R. Franci, 2014. Early adolescent stress alters behavior and the HPA axis response in male and female adult rats: the relevance of the nature and duration of the stressor. Physiol Behav 133(178-189.

Arnsten, A. F., 2009. Stress signalling pathways that impair prefrontal cortex structure and function. Nat Rev Neurosci 10(6), 410–422.

Bandelow, B. and S. Michaelis, 2015. Epidemiology of anxiety disorders in the 21st century. Dialogues Clin Neurosci 17(3), 327–335.

Bangasser, D. A., A. Curtis, B. A. Reyes, T. T. Bethea, I. Parastatidis, H. Ischiropoulos, E. J. Van Bockstaele and R. J. Valentino, 2010. Sex differences in corticotropin-releasing factor receptor signaling and trafficking: potential role in female vulnerability to stress-related psychopathology. Mol Psychiatry 15(9), 877, 896-904.

Bangasser, D. A., K. R. Wiersielis and S. Khantsis, 2016. Sex differences in the locus coeruleus-norepinephrine system and its regulation by stress. Brain Res 1641(Pt B), 177–188.

Bingham, B., K. McFadden, X. Zhang, S. Bhatnagar, S. Beck and R. Valentino, 2011. Early adolescence as a critical window during which social stress distinctly alters behavior and brain norepinephrine activity. Neuropsychopharmacology 36(4), 896–909.

Blume, S. R., M. Padival, J. H. Urban and J. A. Rosenkranz, 2019. Disruptive effects of repeated stress on basolateral amygdala neurons and fear behavior across the estrous cycle in rats. Sci Rep 9(1), 12292.

Boisclair Demarble, J., C. Fortin, B. D’Antono and S. Guay, 2020. Gender Differences in the Prediction of Acute Stress Disorder From Peritraumatic Dissociation and Distress Among Victims of Violent Crimes. J Interpers Violence 35(5-6), 1229–1250.

Borodovitsyna, O., M. D. Flamini and D. J. Chandler, 2018. Acute Stress Persistently Alters Locus Coeruleus Function and Anxiety-like Behavior in Adolescent Rats. Neuroscience 373(7-19.

Borodovitsyna, O., N. Joshi and D. Chandler, 2018. Persistent Stress-Induced Neuroplastic Changes in the Locus Coeruleus/Norepinephrine System. Neural Plast 2018(1892570.

Bradshaw, S. E., K. L. Agster, B. D. Waterhouse and J. A. McGaughy, 2016. Age-related changes in prefrontal norepinephrine transporter density: The basis for improved cognitive flexibility after low doses of atomoxetine in adolescent rats. Brain Res 1641(Pt B), 245–k257.

Curtis, A. L., T. Bethea and R. J. Valentino, 2006. Sexually dimorphic responses of the brain norepinephrine system to stress and corticotropin-releasing factor. Neuropsychopharmacology 31(3), 544–554.

Jedema, H. P. and A. A. Grace, 2004. Corticotropin-releasing hormone directly activates noradrenergic neurons of the locus ceruleus recorded in vitro. J Neurosci 24(43), 9703–9713.

Kessler, R. C., A. Sonnega, E. Bromet, M. Hughes and C. B. Nelson, 1995. Posttraumatic stress disorder in the National Comorbidity Survey. Arch Gen Psychiatry 52(12), 1048–1060.

Kolb, B. and R. Gibb, 2011. Brain plasticity and behaviour in the developing brain. J Can Acad Child Adolesc Psychiatry 20(4), 265–276.

Lovelock, D. F. and T. Deak, 2019. Acute stress imposed during adolescence yields heightened anxiety in Sprague Dawley rats that persists into adulthood: Sex differences and potential involvement of the Medial Amygdala. Brain Res 1723(146392.

McCall, J. G., R. Al-Hasani, E. R. Siuda, D. Y. Hong, A. J. Norris, C. P. Ford and M. R. Bruchas, 2015. CRH Engagement of the Locus Coeruleus Noradrenergic System Mediates Stress-Induced Anxiety. Neuron 87(3), 605–620.

McCall, J. G., E. R. Siuda, D. L. Bhatti, L. A. Lawson, Z. A. McElligott, G. D. Stuber and M. R. Bruchas, 2017. Locus coeruleus to basolateral amygdala noradrenergic projections promote anxiety-like behavior. Elife 6(

McCormick, C. M., I. Z. Mathews, C. Thomas and P. Waters, 2010. Investigations of HPA function and the enduring consequences of stressors in adolescence in animal models. Brain Cogn 72(1), 73–85.

McCormick, C. M., C. Smith and I. Z. Mathews, 2008. Effects of chronic social stress in adolescence on anxiety and neuroendocrine response to mild stress in male and female rats. Behav Brain Res 187(2), 228–238.

Mora, S., N. Dussaubat and G. Diaz-Veliz, 1996. Effects of the estrous cycle and ovarian hormones on behavioral indices of anxiety in female rats. Psychoneuroendocrinology 21(7), 609–620.

Mulvey, B., D. L. Bhatti, S. Gyawali, A. M. Lake, S. Kriaucionis, C. P. Ford, M. R. Bruchas, N. Heintz and J. D. Dougherty, 2018. Molecular and Functional Sex Differences of Noradrenergic Neurons in the Mouse Locus Coeruleus. Cell Rep 23(8), 2225–2235.

Olpe, H. R. and M. W. Steinmann, 1982. Age-related decline in the activity of noradrenergic neurons of the rat locus coeruleus. Brain Res 251(1), 174–176.

Paus, T., M. Keshavan and J. N. Giedd, 2008. Why do many psychiatric disorders emerge during adolescence? Nat Rev Neurosci 9(12), 947–957.

Reyes, B. A., R. J. Valentino, G. Xu and E. J. Van Bockstaele, 2005. Hypothalamic projections to locus coeruleus neurons in rat brain. Eur J Neurosci 22(1), 93–106.

Romeo, R. D., S. J. Lee, N. Chhua, C. R. McPherson and B. S. McEwen, 2004. Testosterone cannot activate an adult-like stress response in prepubertal male rats. Neuroendocrinology 79(3), 125–132.

Romeo, R. D., S. J. Lee and B. S. McEwen, 2004. Differential stress reactivity in intact and ovariectomized prepubertal and adult female rats. Neuroendocrinology 80(6), 387–393.

Selye, H., 1975. Stress and distress. Compr Ther 1(8), 9–13.

Spear, L. P., 2000. The adolescent brain and age-related behavioral manifestations. Neurosci Biobehav Rev 24(4), 417–463.

Toledo-Rodriguez, M. and C. Sandi, 2011. Stress during Adolescence Increases Novelty Seeking and Risk-Taking Behavior in Male and Female Rats. Front Behav Neurosci 5(17.

Valentino, R. J., S. L. Foote and G. Aston-Jones, 1983. Corticotropin-releasing factor activates noradrenergic neurons of the locus coeruleus. Brain Res 270(2), 363–367.

Valentino, R. J., M. Page, E. Van Bockstaele and G. Aston-Jones, 1992. Corticotropin-releasing factor innervation of the locus coeruleus region: distribution of fibers and sources of input. Neuroscience 48(3), 689–705.

van Dijken, H. H., D. C. de Goeij, W. Sutanto, J. Mos, E. R. de Kloet and F. J. Tilders, 1993. Short inescapable stress produces long-lasting changes in the brain-pituitary-adrenal axis of adult male rats. Neuroendocrinology 58(1), 57–64.

Yohn, N. L. and J. A. Blendy, 2017. Adolescent Chronic Unpredictable Stress Exposure Is a Sensitive Window for Long-Term Changes in Adult Behavior in Mice. Neuropsychopharmacology 42(8), 1670–1678.

Zitnik, G. A., A. L. Curtis, S. K. Wood, J. Arner and R. J. Valentino, 2016. Adolescent Social Stress Produces an Enduring Activation of the Rat Locus Coeruleus and Alters its Coherence with the Prefrontal Cortex. Neuropsychopharmacology 41(5), 1376–1385.

